# Development of algorithms for automated detection of cervical pre-cancers with a low-cost, point-of-care, Pocket colposcope

**DOI:** 10.1101/324541

**Authors:** Mercy Nyamewaa Asiedu, Anish Simhal, Usamah Chaudhary, Jenna L. Mueller, Christopher T. Lam, John W. Schmitt, Gino Venegas, Guillermo Sapiro

## Abstract

**Goal:** In this work, we propose methods for (1) automatic feature extraction and classification for acetic acid and Lugol’s iodine cervigrams and (2) methods for combining features/diagnosis of different contrasts in cervigrams for improved performance.

**Methods:** We developed algorithms to pre-process pathology-labeled cervigrams and to extract simple but powerful color and textural-based features. The features were used to train a support vector machine model to classify cervigrams based on corresponding pathology for visual inspection with acetic acid, visual inspection with Lugol’s iodine, and a combination of the two contrasts.

**Results:** The proposed framework achieved a sensitivity, specificity, and accuracy of 81.3%, 78.6%, and 80.0%, respectively when used to distinguish cervical intraepithelial neoplasia (CIN+) relative to normal and benign tissues. This is superior to the average values achieved by three expert physicians on the same data set for discriminating normal/benign cases from CIN+ (77% sensitivity, 51% specificity, 63% accuracy).

**Conclusion:** The results suggest that utilizing simple color- and textural-based features from visual inspection with acetic acid and visual inspection with Lugol’s iodine images may provide unbiased automation of cervigrams.

**Significance:** This would enable automated, expert-level diagnosis of cervical pre-cancer at the point-of-care.

## I. Introduction

### A. Background

In the United States and other high-income countries, cervical cancer incidence and mortality have decreased by 70% over the last 70 years; however, women living in low and middle-income countries (LMICs) still experience a disproportionately high burden of cervical cancer [1]. In fact, about 85% of the global burden of cervical cancer occurs in LMICs, with 87% of cervical cancer related deaths occurring in these regions [2]. In 2012, over 500,000 women were diagnosed with cervical cancer, and approximately 50% of them died from this disease [2]. If status quo is maintained, the numbers are expected to rise to over 750,000 new cases and 400,000 deaths per year by 2035 [2].

In high-resource settings screening for cervical cancer is performed using a multi-tiered paradigm which begins with the Papanicolaou (Pap) smear with human papillomavirus (HPV) co-testing, followed by colposcopy guided biopsy and treatment if needed [3]. Colposcopy uses a low-power microscope to visualize cervix changes that are highlighted with the application of exogenous contrast agents such as acetic acid and, in some cases, Lugol’s iodine. Acetic acid (3% or 5%) application to the cervix causes a reversible coagulation in nuclear proteins and cytokeratin, which primarily affects lesion areas due to their high nuclear protein content [4]. This causes whitening and mosaic textured features in abnormal regions, whereas normal cervix regions remain a light pink color. Normal epithelial cells of the cervix are glycogen rich and take up Lugol’s iodine, which turns them dark brown, while abnormal areas are glycogen deficient and do not uptake the glycophilic Lugol’s iodine solution and hence, appear pale/mustard yellow [4]. Suspicious areas are biopsied for pathology confirmation of cervical abnormalities [3]. Women with pre-cancer are treated via excision of a portion of the cervix using Loop Electrosurgical Excision Procedure (LEEP). Women with cancer are referred to a combination of local and/or systemic therapy depending on the stage of invasive disease. This screening model is not practical to implement in LMICs due to the resources that are required to procure and implement well-established solutions [5, 6].

In low-resource settings, HPV testing has been recommended as an alternative to the Pap smear. However, HPV testing is not widely available in LMICs [5] and, if available, requires a secondary test that provides specificity in order to prevent large numbers of women from being unnecessarily referred for confirmatory testing or treatment [5]. Visual inspection with acetic acid (VIA) using the naked eye is a low-tech version of colposcopy that serves to either add specificity to HPV testing (when available) or serves as the primary screening tool. However, VIA is not scalable for a number of reasons. Specifically, the lack of image capture results in poor quality control due to lack of documentation, and lack of magnification diminishes sensitivity to microscopic disease [7, 8].

Our group has developed a low-cost, portable, point-of-care digital colposcope, called the Pocket colposcope, to replace VIA with high-quality colposcopy, which is the established standard-of-care for triage in high-resource settings [9–11]. An international study conducted by our group found high concordance between the Pocket colposcope and standard-of-care clinical colposcopes [12]. The Pocket colposcope can overcome the limitations of VIA in settings where traditional colposcopes are too cumbersome and/or expensive to use. This will not only provide enhanced visualization and magnification compared to traditional VIA but will also provide much needed documentation and feedback to ensure quality control in healthcare provider performance. However, the limited availability of expert colposcopists in low resource settings represents yet another bottleneck [5, 6]. Therefore, our goal is to develop image processing and machine learning-based approaches for images obtained with the Pocket colposcope to potentially recapitulate the expertise of an experienced colposcopist in settings where there is a shortage of experts available to screen large populations.

### B. Literature review

Recently, deep learning methods have been explored for classifying cervigrams and other cancer imaging applications, such as endoscopy. Xu et al used a multimodal convolutional neural network with inputs from a combination of (1) physician interpretation of 10,000 colposcopy images, and (2) patient clinical records (age and pH value, Pap smear cells, HPV signal and status) to a provide classification model with 88.91% accuracy, 87.83% sensitivity, and 90% specificity [13]. However, limitations include the need for a large data set (10,000 images), and the use of Pap and HPV information, which may not be available, particularly in low-resource settings where incidence and mortality are highest. Additionally, physician diagnosis was used as ground truth, which introduces human subjectivity present in colposcopy. In endoscopy, another internal imaging technique, deep learning has been used for computer aided diagnoses in recognizing and identifying polyps and tumors [14–17].

Even though deep learning has a huge potential application in medical imaging for accurate computer-aided diagnosis (and has seen recent successes in FDA cleared technologies), its application has yet to scale up due to the lack of large datasets comprised of accurately annotated images that transfer across acquisition devices and protocols. Though these can be achieved in cervical colposcopy limitations, addressed here, it will require time, cost, and availability of resources for pathological annotation. Studies have proposed image data generation methods to overcome the obstacle of large data sets needed for deep learning [15, 16]. Though these methods have promise for future applications, they are currently not viable for our application. Finally, domain specific features exploited here are more explainable than those often obtained from deep-learning, and this is critical both for adoption by providers and identifying clinical steps following screening.

Several groups working to create algorithms for automated computer-aided colposcopy have shown proof of concept and feasibility using traditional classification methods with handcrafted feature extraction, classifier training, and validation [18–28]. The majority of these methods have been applied to the use of VIA only, which, on its own, suffers from limitations in sensitivity and specificity [7, 8, 29–33]. Additionally, most of these methods require pre-selection of suspicious regions, and use physician labels for ground truth rather than gold-standard pathology, introducing a level of human error.

Visual inspection with Lugol’s iodine (VILI), which provides another important source of contrast for the visualization of cervical abnormalities, has not been evaluated previously. Studies with physicians providing diagnosis based on both contrasts have shown that VILI has the potential to bolster the performance of algorithms based on VIA alone [10, 34]. To increase specificity and sensitivity of automated diagnosis, our group is developing a method that factors in all sources of contrast physicians consider on the modified Reid Colposcopic Index (RCI) for cervical pre-cancer diagnosis [4].

### C. Study Goal

While some recent tools such as U-NET [35] have demonstrated to be very general for numerous medical imaging tasks, the goal of this work is not to introduce a powerful, generic, new algorithm, but to use domain knowledge to solve a very specific and important challenge: automatic screening of cervical pre-cancers. We demonstrated that with a very simple, computationally efficient, and explainable tool we achieve expert-level diagnosis with limited training data. Some of the lessons learned from the proposed technical contribution, in particular when combined with other more generic tools, could potentially benefit other medical imaging segmentation and screening applications, when properly combined (as done here) with domain knowledge.

We have developed a series of feature extraction and simple machine algorithms that leverage both VIA and VILI images obtained with Pocket colposcope for the automated diagnosis of cervical pre-cancers to recapitulate expert colposcopist performance. By developing a novel strategy incorporating VILI algorithms, we hypothesize we can achieve improved sensitivity and specificity performance over VIA alone. Unlike previous approaches, this method uses pathology gold standard labels for training and does not require a health provider to preselect an area of concern, but rather evaluates the entire cervix to automatically identify regions of interest. The algorithms pre-process images to reduce specular reflection, automatically segments a region of interest from the cervix for analysis, extracts color- and texture-based features, and utilizes a support vector machine for binary classification of VIA and VILI images. Receiver operating characteristics (ROC) curves are generated from the classifier to determine the area under the curve (AUC), which indicates how well a model predicts classes [36]. VIA and VILI algorithms were then combined. With the proposed framework, the best performing model used the combined VIA and VILI and achieved sensitivity of 81.3%, specificity of 78.6%, accuracy of 80.0%, and overall AUCs of 0.86 compared to pathology. As a point of comparison, expert physicians’ interpretation achieved an average sensitivity of 77% and specificity of 51% for the same dataset.

Our main contributions are summarized in the following three areas and their complete integration:

1. An overall real-time, simple, efficient, and repeatable algorithm which utilizes established approaches in image processing and machine learning to classify cervical cancer images. Classification is performed for individual contrasts, and combinations of these contrasts with high accuracy and speed compared to expert colposcopists. We develop algorithms for acetic acid only, Lugol’s iodine only, and different combinations of the two and show a synergistic improvement in performance, compared to using one source of contrast. To the best of our knowledge, this is the first work extracting and combining explainable features from acetic acid and Lugol’s iodine images for cervical cancer classification. The proposed methods for combining contrasts can be potentially applied to other imaging modalities, specifically, clinical colposcopes that use more than one source of contrast for diagnosis.
2. This method is different from other methods for cervical cancer image classification that require preselection of suspicious regions of the cervix, or use physician interpretation as ground truth, which introduces subjectivity and human error. In previous published studies, a region of the cervix is manually selected and then further analyzed for classification. In our study, we automatically segment a region of interest using Gabor filters and then further analyze the segmented regions for color and texture.
3. The Pocket colposcope used in this study is unique compared to other colposcopes in its ability to acquire an image an inch away from the cervix, remove a majority of the noise (speculum, vaginal walls), and consequently decrease the level of image processing complexity required for cervix segmentation, which has been documented in previous methods [9–11]. When combined with the algorithms introduced here, the Pocket colposcope could enable widespread scalable screening for cervical cancer.

## II. Methods

First, VIA and VILI images from the Pocket colposcope were cropped to remove clinically irrelevant features such as the speculum, vaginal walls. Specular reflection was then automatically attenuated to reduce glare artifacts. For both VIA and VILI images, a Gabor filter was applied to automatically segment regions within the cervix for further analysis based on color and texture differences. Haralick’s features (contrast, correlation, energy and homogeneity) [37] were calculated from these segments. The segments were also transformed to different color spaces (grayscale, RGB, YCbCr, HSV and CIElab). From each color space central tendencies of mean, median, mode, variance and Otsu threshold level [38] were calculated for each color channel of each color space. Additionally, for VILI images the percentage of pixels corresponding to the yellow staining from non-uptake of Lugol’s iodine by the lesion region was used to determine a pseudo lesion size. An optimal subset of these features was selected using a wrapper forward sequential feature subselection method with a support vector machine (SVM) for the individual VIA and VILI algorithms. These features were used to train a SVM classifier with pathology as ground truth labels. Five-fold cross validation was used to determine performance and generate ROC curves for binary classification of images as VIA/VILI positive or negative. For the parallel-combined algorithm, scores for classifying the VIA and VILI images with their respective algorithms were input as predictors to generate a classifier for combined classification. For the serial-combined algorithm, the VIA algorithm with selected features was first applied to the VIA image data set at a threshold set to enable high sensitivity which resulted in a high positivity rate. VIA negatives were classified as negatives by the algorithm while VIA positives were further analyzed with the VILI algorithm. Corresponding VILI images of the VIA positives were analyzed with the VILI algorithm (which has higher specificity) for binary classification as negative or positive. Results for VIA only, VILI only, and the combined methods were compared to each other and to the performance of expert physicians on the same data set. Fig 1 provides a summary of the steps of the individual VIA and VILI algorithms described, which is further described below. It is important to note that the algorithm uses a combination of domain-knowledge inspired features with machine learning, since unlike other applications which use neural networks, the amount of training data is limited.

**Fig 1.**
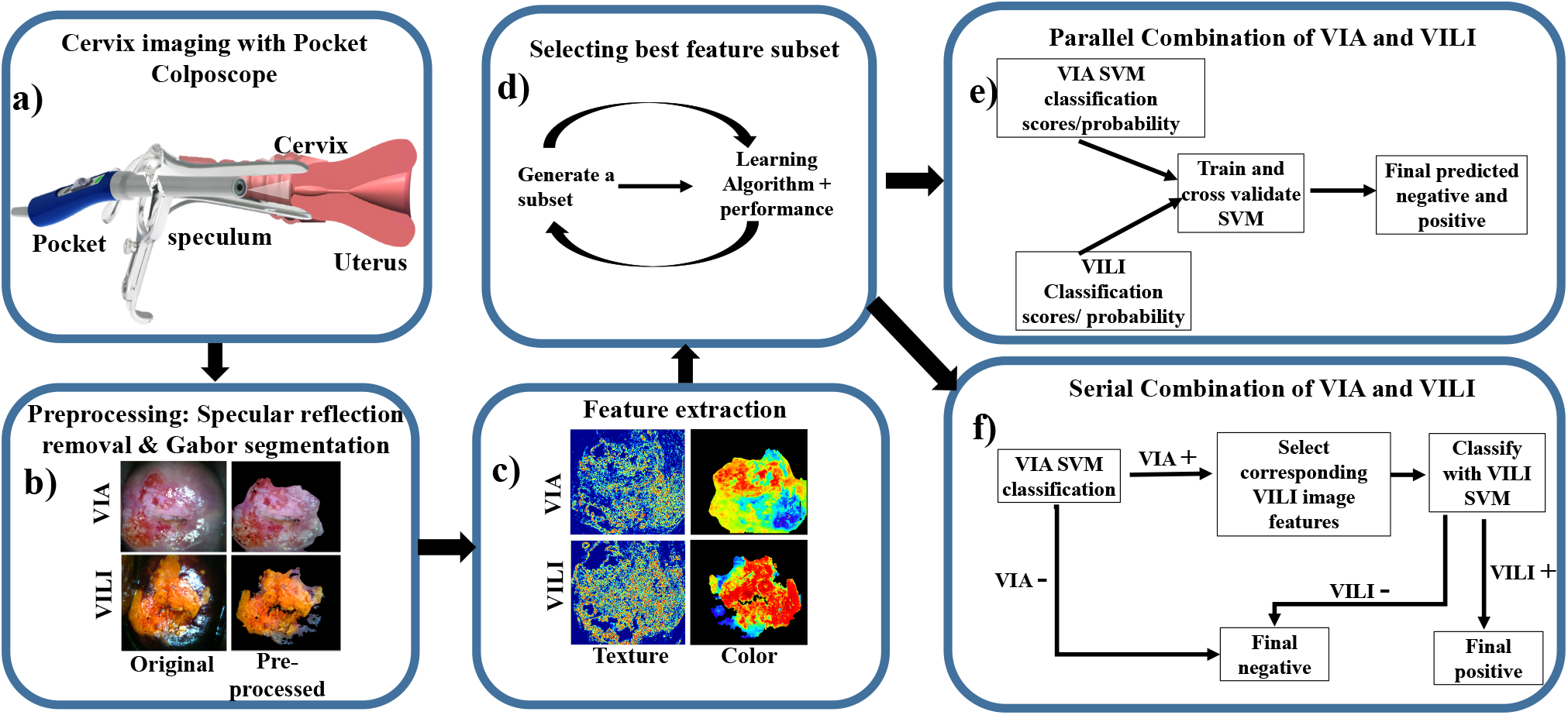
Flow chart of the individual feature extraction and classification process, a) Image collection: A low-cost, Pocket colposcope for cervix imaging was used to obtain 134 image pairs of acetic acid (VIA) and Lugol’s iodine contrasts (VILI), b) Pre-processing: These images were preprocessed by applying a specular reflection attenuation algorithm to remove bright white speckles. Custom Gabor filters were applied to the image to select a region of interest for further processing. The pre-processing and segmentation stages were the same for both VIA and VILI, c) Feature extraction: For VIA pre-processed images, Haralick’s texture features, including contrast, correlation, energy and homogeneity, were obtained. Images were also transformed into individual channels of different colorspaces, overall generating 69 features for each VIA image. For VILI images, the same Haralick’s texture and color features were extracted, however approximate lesion stain size was also determined, yielding 70 features, d) A subset of these features were selected with a wrapper feature selection method and selected features were used to train and cross-validate a support vector machine, (e-f) Finally, the features from VIA and VILI were combined using 2 different methods: e) Parallel method and f) Serial method. These were validated to determine improvement of combined classification over using one source of contrast alone.

### A. Image Collection

Images, pathology and physician labels were retrospectively obtained from a database of Pocket Colposcope images acquired in a previous clinical study [12] in which blinded expert physicians provided a diagnosis for each patient from reviewing randomized digital cervigrams based on both VIA and VILI images.

To summarize, two hundred patients undergoing colposcopy examination at La Liga Contra el Cancer Clinic in Lima, Perú were recruited for a clinical study. This study was approved by Duke University Institutional Review Board (Pro00052865) and performed with approved protocol, informed consent process, and data storage system at La Liga Contra el Cancer, Lima, Peru. For each patient, acetic acid was applied to the cervix and images were captured with the standard-of-care colposcope followed by the Pocket Colposcope. Lugol’s iodine was then applied to the cervix and VILI images were captured with the standard-of-care colposcope followed by the Pocket colposcope. Images, patient demographics, and pathology results were collected and stored in a HIPAA compliant secured database, REDcap [39]. No biopsies were taken for normal colposcopies as per standard-of-care procedures. Hence, the ground truth for VILI negative images was primarily based on expert physician interpretation. Of biopsies taken based on positive colposcopy, 6 came back normal, 19 came back as benign conditions (cervicitis and condilomas) and the remainder were pre-cancers (CIN1, CIN2+) or cancers (invasive carcinomas). For algorithm binary classification labels, condilomas and cervicitis were considered pathologically normal. 25 out of 51 (49%) colposcopy-negatives were pathology-confirmed negative and 26 out of 51 (51%) were negative based on interpretation by the lead colposcopist at the clinic. VIA/VILI image pairs which were deemed interpretable by an expert physician were randomized and sent out to expert colposcopists for cervical pre-cancer diagnosis. Three expert physicians reviewed the images. Two from Duke University Medical Center and one from La Liga Contra el Cancer in Lima, Peru. Experts were blinded to the algorithmic results. Physician interpretation was based on both VIA and VILI features and were classified as either VILI/VIA positive or negative.

### B. Cervix pre-processing

#### Cervix cropping

Cervigrams typically contain clinically superfluous features such as the vaginal sidewalls and the speculum. These artifacts can affect feature extraction and diagnostic accuracy of an automated algorithm and therefore were removed to enable accurate analysis and classification of features pertaining to the cervix. Due to image positioning diversity, the cervix region of interest (ROI) was cropped using a minimum bounding box around the cervix region. With standardized images in which the cervix took up about 90% of the image, no cropping was necessary.

#### Specular reflection attenuation

Specular reflections appear as bright white spots on the cervigram where there is saturation of light exceeding the camera’s linear detection range for a set exposure. Specular reflections from Pocket colposcope images were primarily caused by moisture, the uneven surface of the cervix, excess light illumination, and for the VILI images light reflecting off the dark VILI stains. Specular reflection affects subsequent processing methods, which are primarily color-based, hence the need to attenuate their effect. We employed the well-established specular reflection attenuation method described by Das et al. (2010) [40]. In summary, the raw RGB image was separated into the R, G, and B channels. Specular reflection was identified by finding pixels above a threshold (220 for an 8-bit image) from each channel and logically AND-ing them. In the binary image, the outlines of reflection were dilated, and the borders of dilated outlines were identified on the original image. A Laplacian infill method using values from these borders was used to smoothly interpolate inward from the border pixels [40], providing an output image with reduced specular reflection.

#### Gabor segmentation

A Gabor algorithm combined with k-means clustering was applied to the grayscale of the raw RGB image to select a region within the cervix for further analysis. Gabor filters look for specified frequency content in an image in specified regions and directions, similar to the human visual system [41]. Our hypothesis was that the Gabor filter would have a high frequency response within the acetowhite/texturized regions in VIA positive and yellow lesion regions in VILI positives. K-means clustering, an unsupervised learning method which splits unlabeled data into k number of clusters [42], was then used to segment the Gabor filtered image into two clusters, and the cluster with the higher Gabor mean was selected for further analysis. For VIA/VILI negative regions, the assumption was that a region within the cervix would be selected for further analysis, which would subsequently indicate negativity. Utilizing this approach overcomes limitations from previous studies where regions needed to be manually selected for further textural analysis. A multi-scale Gabor filter was created with different orientations (0, 45, 90 and 135 degrees) and different wavelengths with increasing powers of two starting from 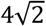 up to the hypothenus length of the input image. The orientations and wavelengths used in this study are well established for unsupervised texture segmentation [41]. The filter was convolved with each image to obtain the overall Gabor response. K-means clustering (k=2) was used to segment the cervix into two regions based on the Gabor response. The region with the highest mean Gabor response was selected for further analysis. Gabor segmented regions were used to calculate color, and Haralick’s texture features for both VIA and VILI images. Pseudo lesion size was also calculated for VILI images.

### C. Feature extraction

#### Haralick’s texture features

Since acetic acid application causes textured mosaicism, which highlights different vasculature patterns corresponding to different diagnosis, we calculated Haralick’s texture features [37] to analyze the Gabor-segmented VIA regions. This method was also applied to VILI images with the assumption that since VILI is applied after VIA, there may be texturized regions within the VILI segments. In the greyscale image, we calculated the grey-level cooccurrence matrix (GLCM), a second order statistical method which characterizes an image by calculating how often a pair of pixels with specific values and spatial relationship occur in an image. The GLCM was computed for four different pixel offsets (1, 5, 10 and 15) each in four different directions (0, 45, 90 and 135 degrees), for a total of 16 matrices. Small to medium offset values were used since we expected mosaicism features to show pixel differences within this range. These different directions were used since features of interest may be horizontal, vertical or diagonal. From these GLCMs, four Haralick’s features were calculated: contrast, correlation, energy and homogeneity were calculated. This yielded 4 texture-based features for each VIA and VILI image.

#### Color space transformation calculations

According to the mRCI, both VIA and VILI cervigram classification are primarily based on staining. For VIA, acetowhitened regions correspond to lesions, while light pink regions correspond to normal epithelium. For VILI, yellow regions correspond to lesions, while darker brown staining corresponds to normal epithelium. Based on this biological effect explained in detail in the introduction, color-based features were extracted. Each image was transformed to five different color spaces, and these were separated into individual color channels, specifically ‘Red(R), Green(G), Blue(B) (RGB)’, ‘grayscale’, ‘Hue(H), Saturation(S), Value(V) (HSV)’, ‘Luminance(Y), blue chroma(Cb) and Red chroma(Cr) (YCbCr)’, and ‘Lightness(L), red/green(a), and yellow/blue(b) (CIELAB)’ [43]. RGB color space is widely used for digital image acquisition; however, one of its primary disadvantages is that it is device dependent and sensitive to illumination differences, which limits the use for color-based image analysis [44]. CIELAB color space extends the RGB color space from 90% to 100% of all perceivable colors and enables more quantitative applications since it correlates to perceptual descriptors of color [45]. Additionally, CIELAB and the YCbCr color spaces separate image luminous intensity from color information, reducing factors resulting from how an image was taken, such as variations in LED brightness [43]. HSV rearranges the geometry of RGB color space to make it more intuitive and perceptually relevant, separating color based of its hue, saturation and lightness [43]. Gray scale of an RGB image has luminance information [43].

For each color channel, we determined central tendencies of mean, median, mode, and variance. Otsu threshold level (two-class classification) was also determined on color-space transformed images [46] to take advantage of multiple clusters present from the brown normal regions and yellow abnormal regions. The Otsu threshold is a widely used, nonparametric, unsupervised method, which automatically selects a threshold to separate an image into two clusters, maximizing inter-class variance and minimizing intra-class variance [38]. From each of the thirteen channels of the different color space transforms, the mean, median, mode, variance, and Otsu’s threshold were extracted as potential classification features. Overall 65 color-based features were extracted for each VIA and VILI image.

#### Lesion size determination

A pseudo lesion size detection was performed for VILI images after Gabor segmentation. Due to the high variation in acetowhitening lesion and normal epithelium shades, it was not possible to determine lesion size for VIA images based on acetowhitening alone. Since VILI images provide a higher, more consistent contrast between dark brown normal epithelium and yellow stained abnormal epithelium, color-based “pseudo lesion size” was determined using VILI images. VILI images were transformed into CIElab color space explained previously and the b channel was extracted due to its direct linearity of yellow shades and high contrast in color transforms. A threshold was determined via analysis of pixel intensities of ROIs from a subset of images corresponding to lesions and non-lesion regions. The number of pixels above this threshold was divided by the total number of pixels in the bounded cervix region to approximate a percent “lesion size”. We call it pseudo lesion size because Lugol’s iodine solution may also result in yellowing due to non-uptake in benign cervical conditions, such as condilomas, cervicitis. Parts of the cervix where Lugol’s Iodine is not properly applied (such as cervix boundaries) also show non-uptake yellow features. This resulted in overall a total of 70 features for VILI and 69 features for VIA images.

### D. Feature selection, classifier training and testing

#### Classifier design

We selected a support vector machine classifier, a widely used and successful supervised learning method, which outputs an optimal hyperplane for classification [43, 47, 48]. SVM parameters are as follows:

*Predictor standardization*: Predictors standardized using their corresponding weighted means and weighted standard deviations. *Kernel function*: Gaussian or RBF.

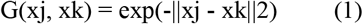

*Kernel Scale*: ~1.008 by automatic appropriate scale factor determined using a heuristic procedure. This heuristic procedure uses subsampling, so estimates can vary from one call to another. *Solver*: ISDA Iterative Single Data Algorithm *Kernel offset*: 0.1

A labeled training data set is input into the classifier which then builds a model to enable classification of a new data set. For non-linearly separable data sets, the SVM can transform data points to higher dimensions to linearize them for optimal separation using higher order kernels [49]. From initial scatter plots demonstrating non-linearity of the data set, we selected a Radial Basis Function (RBF or Gaussian) kernel with automatically selected kernel size using cross validation. RBF kernel was selected because it is most widely used due to flexibility and ability to project data into high (infinite) dimensionality space to find a linear separation [49]. The features were used as predictors while the pathology diagnosis (normal and benign vs. CIN+) was used as input labels for binary classification. While other machine learning tools could potentially be used, SVM was selected due to the limited data available for training and its excellent performance.

#### Feature selection and validation

In machine learning, classifying data using a proportionally high number of features compared to sample size can result in classifier failure due to redundancy and overfitting or overtraining to noise/artifacts. High feature number also increases storage and computational requirements. Due to a limited sample size (n=134 VIA/VILI pairs), a small subset of important features for satisfactory classification performance was selected using forward sequential feature selection (FSFS) with minimum classification error (MCE) [50]. The FSFS is a fast, wrapper feature selection method, similar to the SVM recursive feature elimination (SVM-RFE) method. In the FSFS, feature candidates were sequentially added to an empty set to create a subset, until there was no further improvement in prediction. For each feature subset, the function performed a 5-fold cross validation with the classifier. The forward sequential feature sub selection method is advantageous because unlike filter methods which simply look at performance of individual features, FSFS considers performance of features when combined, as well as feature correlation. The feature number and combination, which provided the minimum error for classification was then selected.

#### Training and testing

The optimal feature subset was used as predictors into the SVM to generate a classification model which was validated using 5-fold cross validation with pathology as ground truth (note that pathology is not available even to clinicians doing visual observation, so this ground truth is a very high standard). For cross validation for VIA and VILI features, the data set was divided into fifths. Training was performed on 4/5 of the data and testing performed on the remaining 1/5 of the data. This was repeated leaving a different data set each time and the results averaged. ROCs were generated to determine the AUC, and sensitivity and specificity of each the algorithm was compared to physician performance for classifying images as negative or positive.

### E. Combining VIA and VILI features

#### Parallel combination

When the SVM classifier classifies VIA and VILI images as negative and positive it also assigns a score to each image which describes the probability or likelihood that the label comes from one of the classes. These probabilities for VIA and VILI images were used as predictor inputs into the parallel-combined SVM classification model with binary pathology labels. A classification model was then generated from these to provide binary classification based on both VIA and VILI inputs.

#### Serial combination

For serially combining VIA and VILI results, we analyzed the data from the individual VIA and VILI algorithms to determine where they complemented each other. Based on this analysis, we first ran the VIA algorithm, using the same feature subset on all images with a low classification cutoff (−0.25 instead of 0) for binary classification. This enabled classification of all high grades but also resulted in a high false-positivity rate. Negatives based on this classification were considered true negatives. The VILI algorithm was then run on the VIA positive images to increase specificity. An SVM classifier model was generated for the VIA and then VILI stage and cross-validated with pathology as ground truth.

## III. Results

### A. Image breakdown

Of the 200 Pocket colposcope patients, 66 were excluded due to one or more of the following factors: screen fail (2.5%), missing pathology for ground truth (2%), images not saved (missing) (6%), blurriness (10%), and device set at the wrong working distance (7%). Device related issues have been addressed in a newer version of the Pocket colposcope design and improved control software. For the purpose of this study, data from a total of 134 patients were used for image analysis. The pathology distribution of patient whose cervigrams were utilized for the study are outlined in Table 1.

**TABLE I.**
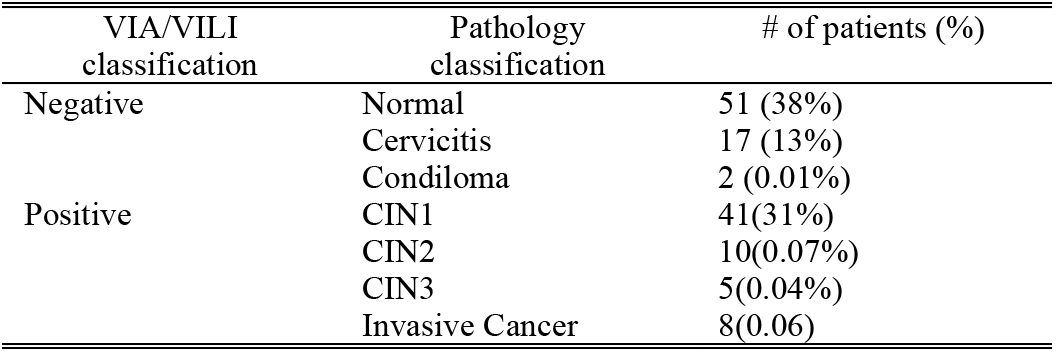
Image Pathology Distribution (N=134)

### B. Image processing and feature extraction Specular reflection attenuation

Pre-processing to attenuate specular reflection was performed prior to feature extraction and classification to prevent inaccuracies due to glare. Fig 2 shows results from representative images with specular reflection before and after specular reflection attenuation. Due to the speckled nature of specular reflection it is difficult to accurately hand select all glare regions for quantitative comparison and analysis of reduction. However, upon visual inspection all images had significantly reduced specular reflection without affecting contrast in the images. This finding is similar to previous work in this area [40].

**Fig 2.**
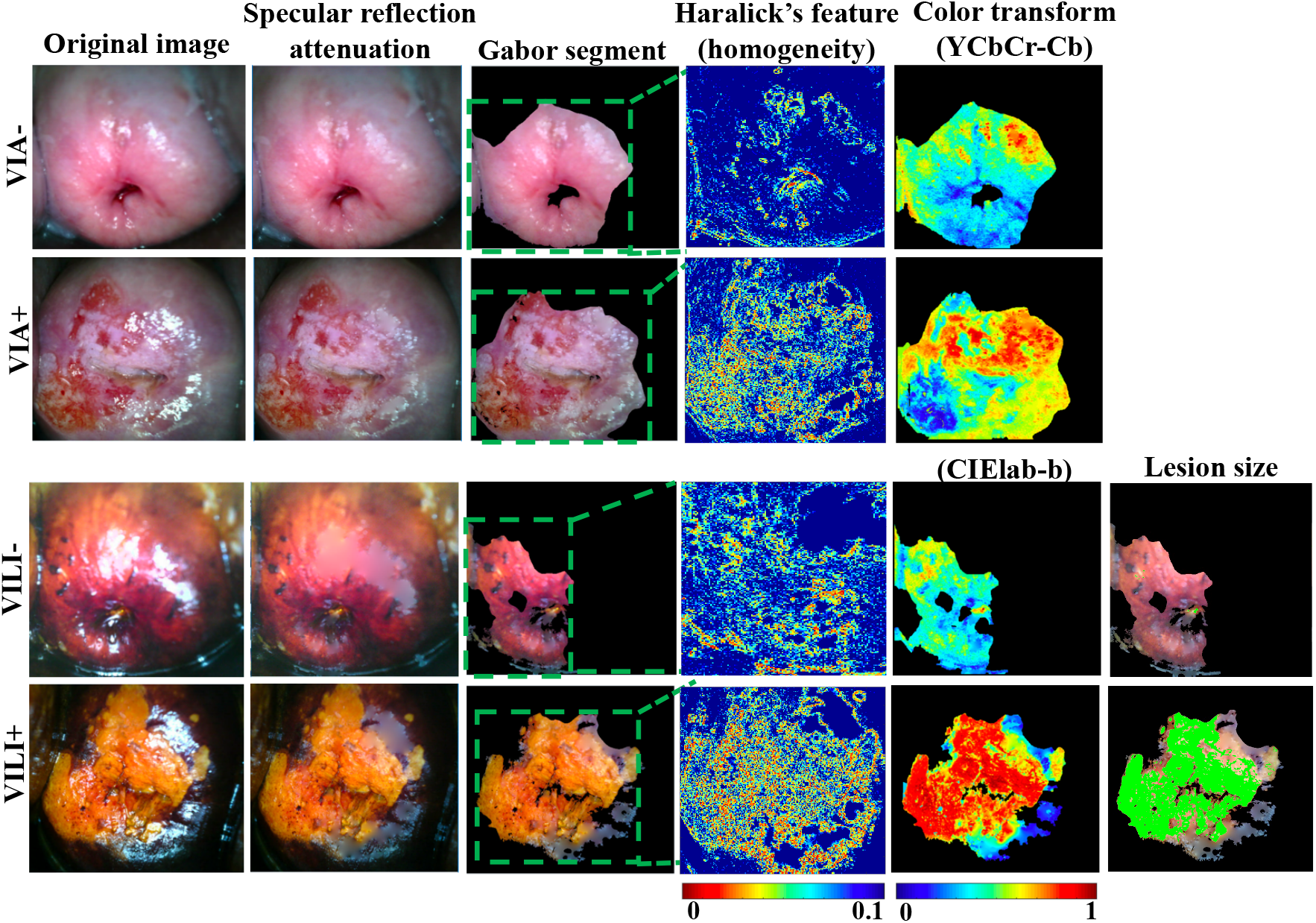
Endpoints for feature extraction methods used for VIA and VILI classification. From left, figure shows original VIA negative and positive images, and VILI negative and positive images. This is followed by specular reflection attenuation results which demonstrate significantly reduced speckle and convincing infilling. Gabor segmentation selects a highly texturized region of the image for further processing. The area around the os is selected and for positive regions in particular, the area around the lesion. Haralick’s texture output of the image is also shown with low homogeneity in lesion regions and high homogeneity in non-lesion regions. Color transforms of the segments are also performed to take advantage in staining differences for negative and positive cervix images. Finally, for VILI images, approximate lesion size is determined.

A Gabor filter was applied to pre-processed VIA/VILI images and the response normalized. Lesion regions for both VIA and VILI Images had higher Gabor responses than the surrounding epithelium. Images were segmented into two main regions by k-means clustering (k=2) and the region with highest mean Gabor response selected. Since VIA and VILI negative images had no or little acetowhitening/yellowing, regions selected were likely to be a subsection of the normal epithelium of the cervix, the os, or the transformation zone. From direct observation, the Gabor segments were in approximately the same regions for both VIA and VILI for 130/134 of the images. The Gabor segmented enabled the automatic selection of a region of interest for further analysis with the color and Haralick’s texture features.

#### Haralick’s texture features

VIA positive images had significantly higher contrast (p=0.03), not significantly different correlation, significantly lower energy (p=0.013), and significantly lower homogenity (p=0.02) compared to VIA negative images. This is expected since “contrast” corresponds to variations between pixels (0 for a constant image), correlation looks at a mutual relationship between pixels (range: −1 to 1, with Nan for a constant image), energy crresponds to uniformity, with higher values for similar pizels (0-1 with 1 for a constant image) and homogenity reflects closeness in distribution to the diagonal of GLCM elements (range: 0-1, with 1 for diagonal/constant image), which calculate how often a pair of pixels with specific values and spatial relationship occur in an image. VILI positive images had significantly higher contrast (p=0.003), insignificantly different correlation, significantly lower energy (p=0.024) and significantly lower homogeneity (0.015) compared to VILI negative images.

#### Color transforms features

Each image was transformed into 5 main color spaces and the central tendencies and Otsu threshold calculated. Fig 2 shows representative color transforms for the VIA (YCbCr-Cb) and VILI (CIElab-b) images. As shown, lesion regions for VILI tended to have lower values than non-lesion regions in the b channel. In the Cb channel for VIA, lesion regions tended to have higher values than non-lesion regions. Different color channels showed distinct trends as well as redundancies. The optimal combination of features was later addressed using the sequential feature selection method. Representative images for all color channels are shown in the appendix.

#### Pseudo lesion size detection

Pseudo lesion size detection was performed on pre-processed VILI images since they enabled higher color-based contrast between lesion and nonlesion regions than VIA. We found that a threshold of 40 in the b channel of CIElab color space enabled adequate cutoff between the yellow and brown regions of the VILI images, the percentage of pixels above a threshold of 40 (which corresponds to all shades of yellow) was divided by the total cervical area to compute lesion size. The mean lesion size was found to be significantly larger (p=0.0047) in VILI positives than in VILI negatives as expected.

### C. Feature selection

Feature selection was performed to select a subset of features to prevent overfitting. Rather than use a simple filter method which selects features with the highest p-values, we used a wrapper forward sequential feature selection (FSFS) method which takes into consideration feature redundancies and interaction. The list of selected features and their p-values are in Table II. Fig 3 compares results of classification scores for VIA and VILI, achieved using features selected from the simple filter selection method and the forward sequential feature selection (FSFS). Results from VIA and VILI images show that features selected with the FSFS method shows expected trends in pre-cancer grades and significantly higher differences and AUC between normal/benign and CIN+ compared to intuitively selected features from the simple filter selection method.

**TABLE II.** Features and P-values From Simple Filter Method and Sequential Feature Sub-selection Method.

| VIA |  |  |  | VILI |  |  |  |
| --- | --- | --- | --- | --- | --- | --- | --- |
| Features w/top p-values |  | Sequential selected |  | Features w/top p-values |  | Sequential selected |  |
| Feature | P-value | Feature | P-value | Feature | P-value | Feature | P-value |
| Correlation | 0.0009 | A mode | 0.07108 | CB median | 0.0001 | B mode | 0.0005 |
| Blue variance | 0.0016 | B mode | 0.2077 | CB mean | 0.0001 | Red variance | 0.5681 |
| L variance | 0.0065 | CR mode | 0.6073 | CB mode | 0.0001 | CB mean | 0.0001 |
| Y variance | 0.007 | Energy | 0.0009 | CB level | 0.000195 | CB level | 0.0002 |
| Grey variance | 0.007 | Correlation | 0.90849 | H mean | 0.00024 | Red mode | 0.17415 |
| Green variance | 0.0147 | CR level | 0.5717 | B mean | 0.00042 | V mode | 0.3329 |

**Fig 3.**
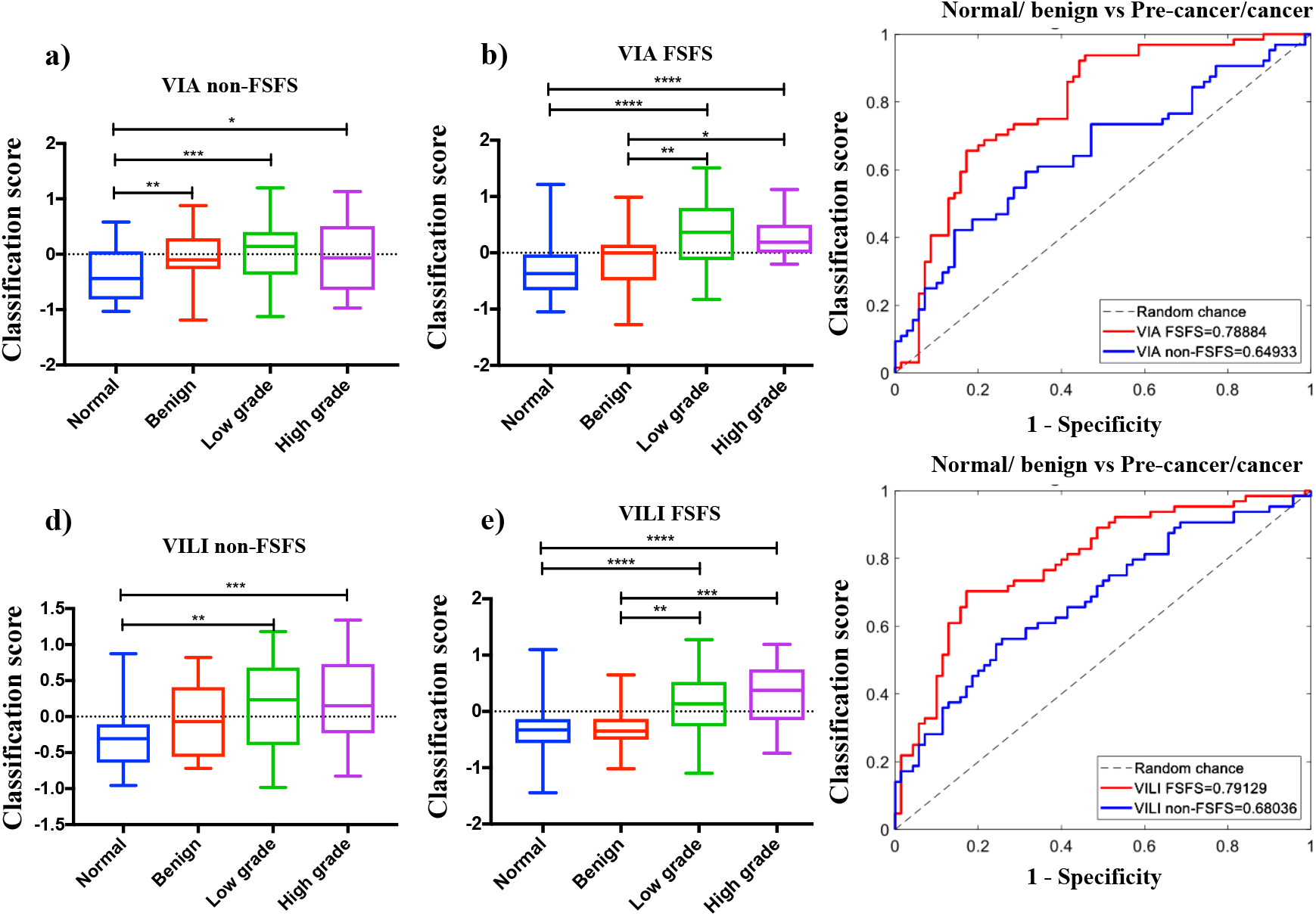
Comparing classifications scores from using top features by p-value (simple filter method) and by FSFS method. a) Box plots of classification scores for VIA algorithm using the features with the highest p-values, b) Box plots of classification scores for VIA algorithm using features selected with sequential feature sub-selection method, c) ROC curves comparing VIA results from features with simple filter method to FSFS method, d) Box plots of classification scores for VILI algorithm using the features with the highest p-values, e) Box plots of classification scores for VILI algorithm using features selected with sequential feature sub-selection method, f) ROC curves comparing VILI results from features with simple filter method to FSFS method. *=p<0.05, **=p<0.001, ***=p<0.0001, ****=p<0.00001.

### D. SVM Classification

Fig 4a and b show scatterplots of classification scores for the parallel and serial combination methods. Classification scores are outputs from the SVM classifier indicating the probability/likelihood of an image belonging to either the negative or positive class. Scatterplots show similar expected trends for both methods. ROCs were generated to compare how the different algorithms (VIA, VILI parallel combined and ground truth for normal/benign vs. CIN+, normal/benign vs. low grade/CIN1 and normal/benign vs. high grade/CIN2+. Performance of three expert physicians (>15 years of colposcopy experience per person) for the same data set are also indicated on the curve. The physician interpretations were collected retrospectively and were based on features from both VIA and VILI image interpretation. Hence physicians had the same information as the combined VIA and VILI algorithms. All physician performance points fall within the ROC curve for combined VIA and VILI algorithms, indicating superior algorithm performance, which is critical considering experts like the ones used in this study are not widely available and certainly lacking in low resource areas. Both combined algorithms outperformed the algorithms using the single contrasts but achieved similar AUCs. Additionally, AUCs for VIA and VILI only AUCs were also similar. Fig 5 shows algorithm and each individual physician’s sensitivities, specificities and accuracy as compared to pathology (Fig 5a and b). Sensitivity (also known as the true positive rate) measures the proportion of correctly identified positive. Specificity (also correctly identified negative cases. Accuracy measures the overall proportion of correctly identified cases. Overall accuracy for all algorithms is higher than physician accuracies. Though physician’s sensitives are on average on par with algorithms, algorithms achieve higher specificities. Combined algorithms achieved highest agreement with physicians, particularly with physicians who had higher performance with pathology. This is predictable since physicians also used both VIA and VILI images for prediction. Fig 5c shows the percentage of correctly classified images by pre-cancer grade for one of the combined (serial) algorithms and the best physician. The algorithm outperforms the best physician across all cervix types but most notably for benign lesions which tend to be overcalled by physicians. Fig 5d-k shows representative VIA/VILI cervigrams of true positive, true negatives, false positives and false negative images classified by the serially combined algorithm. Fig 5d-g show VIA images with their corresponding VILI cervigrams in Fig 5h-k. True positive images showed acetowhitening and mosaicism in VIA image (Fig 5d) and yellow staining in VILI image (Fig 5h). True negatives showed little to no visible whitening or mosaicism in VIA positive image (Fig 5e) and showed brown staining from iodine uptake in VILI image (Fig 5i). False positive images classified as negative by pathology but positive by the algorithm were due to high texture in the VIA image (Fig 5f) and the non-uptake of Lugol’s iodine in the VILI image (Fig 5j). The false negative showed no distinct acetowhitening in the VIA image (Fig 5g) and very little area of Lugol’s non-uptake in the VILI image (Fig 5k). This may be a result of the limitation where biopsies are taken from the endocervical canal which is not typically visible in the image. To conclude, these images as well as all the errors of the automatic algorithms “make sense” and are a result from non-trivial images. Recall that we compared with pathology, which contains data not available to the algorithm or to the clinician.

**Fig 4.**
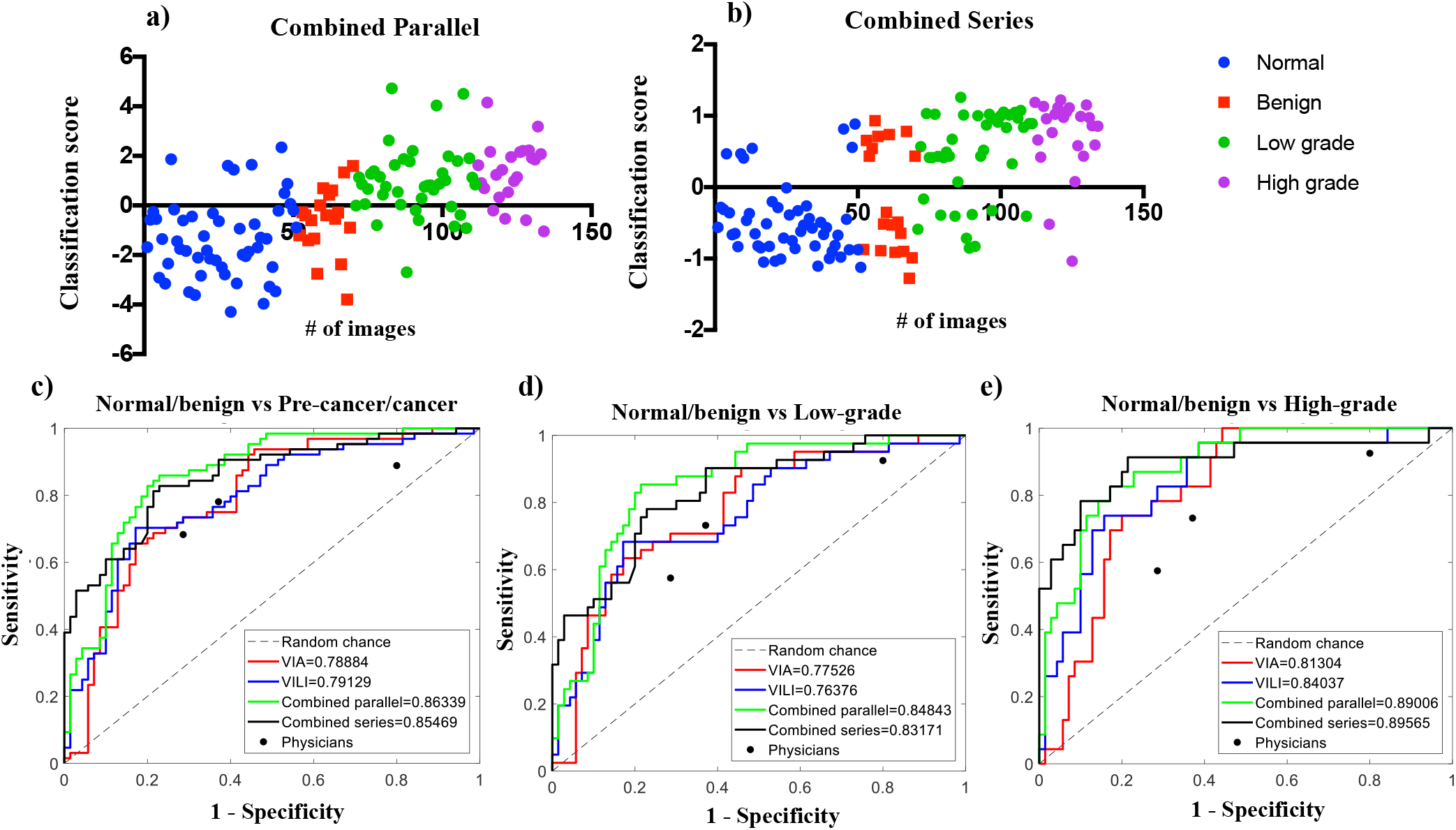
Performance of different classification algorithms, a) Scatterplotfor classification scores from parallel combined method, b) Scatterplot for classification scores from serial combined method, c) ROC curves for all algorithms for normal/benign vs CIN+, d) ROC curves for all algorithms for normal/benign vs low grade/CINl, e) ROC curves for all algorithms for normal/benign vs CIN2+. Performance of each expert physician reader (n=3) is also indicated for each graph. # of images refers to the number of images/ image number.

**Fig 5.**
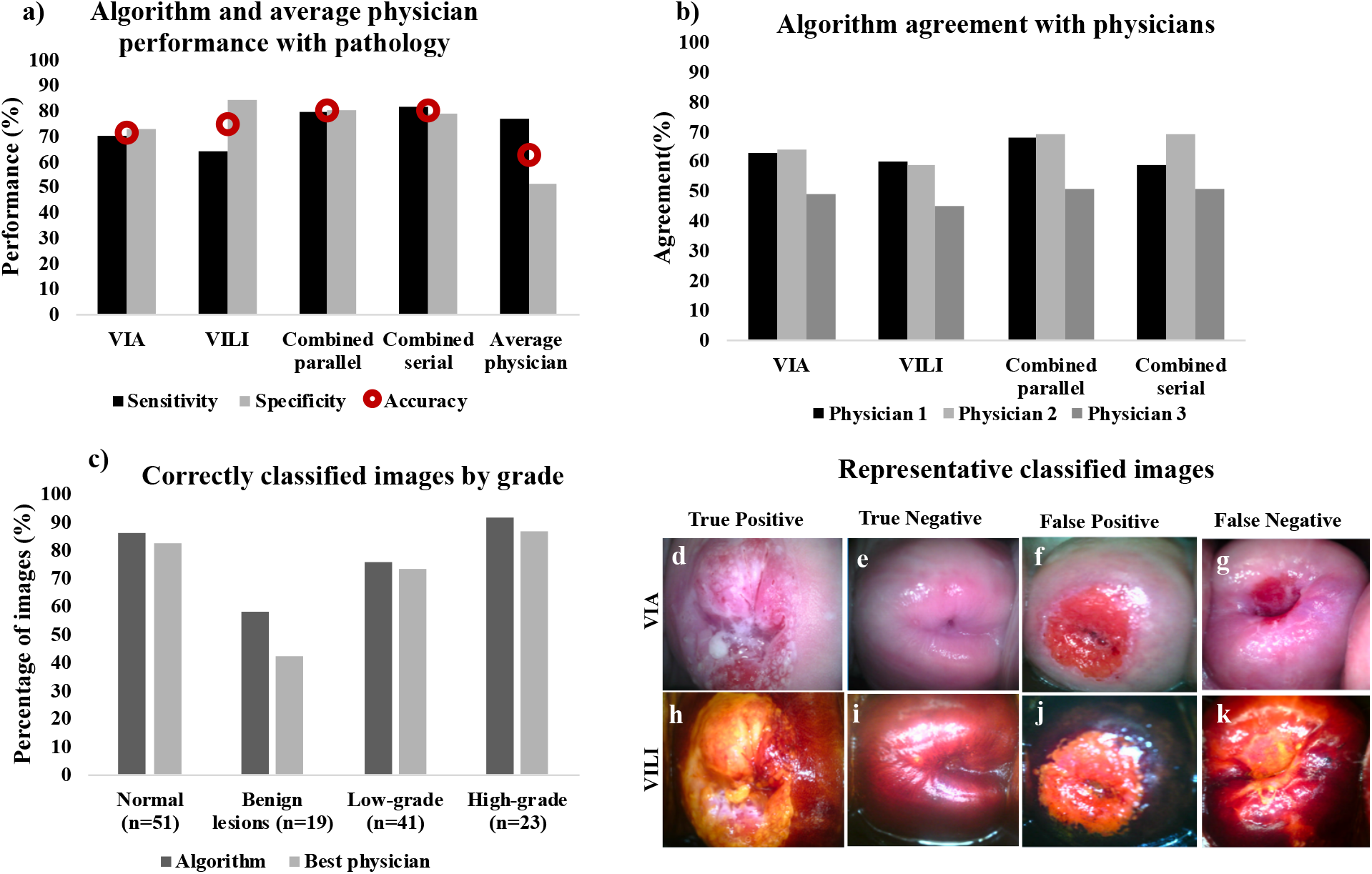
a) Bar charts showing sensitivity, specificity and accuracy (red circular marker) with gold standard pathology for the 4 algorithms; VIA only, VILI only, combined parallel and combined serial compared to average physician performance, b) Bar charts showing percent agreement for each of the algorithms compared to the 3 individual physicians, i.e by what percentage each algorithm agreed with each physician, c) Bar charts showing number of correctly classified images by pathology for the best performing algorithm and the best physician, (d-g) Representative VIA images (h-k) corresponding VILI images diagnosed by algorithm: (d, h) True positive, (e, i) True negative, (f, j) False positive and, (g, k) False negative.

### E) Processing Time Analysis

We ran 10 randomly selected images through the algorithm and measured the machine time taken for individual steps and the overall time. For each image pre-processing for specular reflection attenuation and Gabor segmentation took the most time, averaging 27.7 seconds and 21.6 seconds, respectively, while feature extraction and classification took less than a second. Overall time taken per image averaged 50.3 seconds, and was 2 minutes at most, demonstrating the feasibility for real time diagnosis of images.

**Fig 6.**
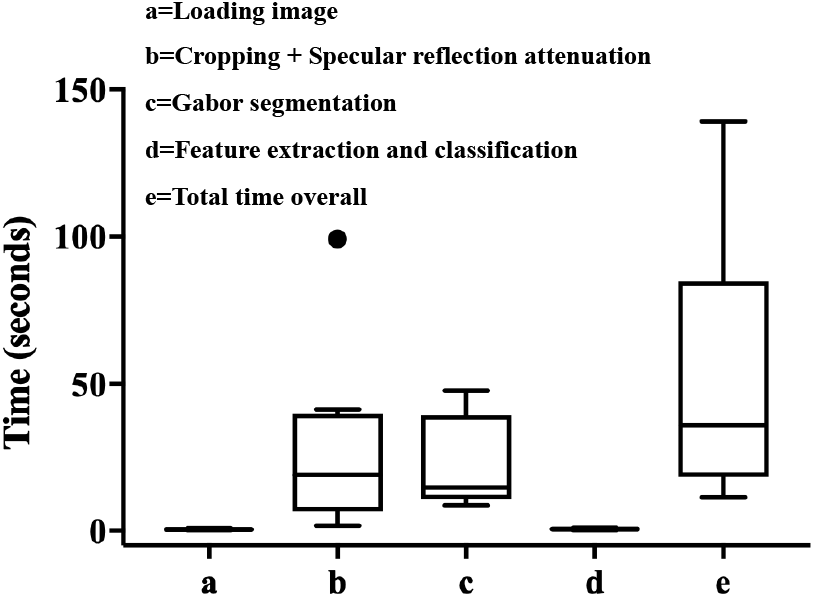
Box plots showing time taken (in seconds) to load, pre-process, process and classify a sample of the images (n=10) as positive or negative for precancer. All but one of the images had a processing time of less than 2 minutes.

## IV. Discussion

We introduced in this study algorithms that process and classify, in real time, VIA and VILI cervigrams as either negative or positive for cervical pre-cancer, and combine features from both contrasts to improve overall specificity while maintaining sensitivity. Algorithms for the individual contrasts were developed to extract color and texture-based features.

A subset of the domain/expertise-motivated features was automatically selected and used to train a support vector machine to develop models for binary classification of cervigrams. The individual algorithms were combined both in parallel (simultaneously) and serially (VIA followed by VILI) to improve performance. Combined methods performed on par with each other, with the parallel method achieving a sensitivity, specificity, accuracy and AUC of 79.7%, 80.0%, 80.0%, and 0.86, respectively, and the serially combined algorithm achieving a sensitivity, specificity, accuracy, and AUC of 81.3%, 78.6%, 80.0%, and 0.86, respectively. This was higher than the average expert physician, who achieved a sensitivity, specificity and accuracy of 77%, 51%, and 63%, respectively. The algorithm more accurately classified negatives, benign conditions, low grades, and high-grade conditions than the best performing physician.

This algorithm has been developed with images captured using the Pocket colposcope, a low-cost, portable, digital colposcope. The labeled images used for this study were obtained retrospectively and physicians used both VIA and VILI data, hence we were not able to directly compare the individual algorithms for VIA and VILI. No previous algorithms for combining VIA and VILI classification have been developed to which we can compare our results. There was only one study we found that had proposed a method to combine VIA and VILI images for diagnoses [51]. However, though preliminary representative images from color-based segmentation were shown, no quantitative performance of the VILI algorithm was provided to which we could compare our results. Additionally, there is no previous algorithm that uses VILI alone for cervical pre-cancer diagnosis. Several groups have proposed methods towards automatic VIA diagnosis which, include automated cervix segmentation, specular reflection removal, and acetowhitening detection using color-based and texture features [25, 52–55]. However, most of these are only semi-automated, requiring manual ROI selection and focusing on lesion detection within abnormal VIA images. Manually selected regions are then classified as a type of cervix tissue using various classifiers. Even though some groups have started working to enable full automation, the current methods cite device- and illumination-dependence, and image diversity being cited as a major challenge in analysis.

Physician performance in our study is on par to those from previous studies. A colposcopy study by Qureshi et al. with 328 women found a sensitivity of 86.84% and a specificity of 48.93f% [56]. A study in Tanzania by Ngoma et al of 10,367 women screened with VILI found a range of 79.8% - 99.3% for sensitivity and 97.0% - 97.6% for specificity [57]. Sankaranarayanan et al performed a study with 4,444 women in Kerala, India, and found a range of 80.6% - 92.0% for sensitivity and 83.6% - 85.8% for specificity [58]. Our physician performance ranged from 68.3 - 88.9% for sensitivity, 20.0 - 71.4% for specificity, and 52.6 - 70.1% for accuracy. It is interesting to note while the average sensitivity of physicians in this study was comparable to that reported in previous studies, the average specificity was lower. We speculate that this may be because physicians who participated in our study were viewing still images of VIA/VILI without any information on patient history or demographics compared to previous studies in which images were interpreted in real time. Additionally, the prevalence of disease is lower at the population level, which was the study population used in published studies, compared to the diagnostic population (secondary screening), which was the study population investigated here. In the diagnostic population, there is a higher number of borderline benign conditions, which can be easily misinterpreted due to intrinsic properties of the contrast agents. Future studies will include testing the algorithm in a real time clinical setting and comparing its performance to pathology and expert physician interpretation in the field.

There are a few limitations related to this work. The first is that this methodology does not account for noisy images. Though there is an extensive training package with the Pocket colposcope on how to obtain clear images, we acknowledge that, with human error, clear images may not always be obtained. Other papers have established methods for recognizing and indicating noisy images. Future renditions of the algorithm will seek to optimize and incorporate these algorithms, such that during a use case, the health provider will be alerted when an image fails to meet quality requirements. An additional limitation is that, even though pathology labels used in this work are considered the clinical standard, there is a degree of subjectivity when a pathologist is histologically grading cervical carcinomas. However, pathological diagnosis has a lower degree of subjectivity compared to physician diagnosis from colposcopy images. Thirdly, even though most of the algorithm is automated, there is still a need to crop the cervix to find a rough ROI in the image using a simple, and easy to use square cropping tool (the only manual feature involved in processing). We have developed a cervix segmentation tool for full automation, but it currently suffers from lack of image standardization due to different working distances when data is captured. In the short term we will incorporate standardization measures into the Pocket colposcope such that images are taken at about the same working distance. This will enable us to apply the segmentation algorithm efficiently, thus fully automating the process. Long term, as we acquire more data with the Pocket colposcope we anticipate using successful deep learning methods such as U-NET, proposed for biomedical image segmentation, to perform this step.

Lastly, classification with this methodology is based on handcrafted color and textural features, identified by expert physicians as predictive of cervical pre-cancer and outlined by cervical cancer guidelines. There may be additional, intrinsic features available in the images, not recognized by the human eye, which can be assessed using deep learning methods. However, due to the limited nature of our data, we are unable to implement these methods and transfer learning efforts have failed to yield accurate results.

While tools from deep-learning have become state-of-the-art in many disciplines, including in medical imaging applications (different than the one addressed here [35, 59]), we should note that they often need significant training data, orders of magnitude beyond what was used in this work. Even when exploiting deep-learning, combined with transfer and/or domain learning to compensate for reduced training data, when the algorithms are deployed they tend to suffer from two fundamental caveats: lack of full explainability AND computational complexity (which translates into energy consumption in mobile devices, critical for low-resource areas). As demonstrated in this paper, they are not actually needed, and through use of domain knowledge we can achieve excellent results with limited training data and with an algorithm at very low computational cost. Comparing results to a previously published study by Xu et al. (2016), which used deep learning methods, with multimodal convolutional neural networks and input of a combination of 10,000 colposcope images (about 2 orders of magnitude more than in this project) and patient clinical records (age and pH value, Pap smear results, HPV signal and status), yielded 88.91% accuracy, 87.83% sensitivity, and 90% specificity [13]. However, Pap and HPV combinations are not readily available in low-resource settings where cervical cancer incidence and mortality is highest, so the real life application of this method is unclear. Visual inspection with acetic acid is the standard of care, and the Pocket colposcope provides the ability to bring high level cervix image capture (colposcopy) to low-resource settings. We present a method which uses a fraction of the data with simple, but powerful features from Pocket colposcopy images to achieve performance of 80% accuracy, 81.3% sensitivity, and 78.6% specificity, comparable to that expert physicians and the deep learning method described above. However, as the amount of data increases with further implementation of the Pocket colposcope, our algorithms, specifically the segmentation stage, could benefit from the U-Net deep-learning mentioned above, which will be the subject of future research.

Long term, as we continue to accumulate data with various clinical studies using the Pocket colposcope, we foresee incorporating deep learning methods to further enhance the hand-crafted, domain-expert features. Given a substantial initial sample size (order of 1000s) from longitudinal clinical use of the Pocket colposcope, we plan to employ data augmentation techniques such as transfer learning, patches, and transformations to increase the size of our dataset. This will enable us to leverage successful deep learning architectures such as convolutional neural networks to extract latent features without the need for image pre-processing steps such as cervix segmentation. Features can then be used in a traditional classification model like logistic regression or support vector machines and thus further improve our classification performance.

Overall, the algorithm for automated cervical cancer diagnosis performed significantly better than average diagnosis of expert physicians. Even though this algorithm is developed using images captured with the Pocket colposcope, the method can potentially be applied to cervigrams from other colposcopes. Overall, this method has potential to enable colposcopy diagnosis in the absence of an expert physician or to complement his/her judgment. Combining VIA and VILI classification outperforms physicians, thus allowing accurate referral colposcopy to be more accessible at the primary care/community level setting.

## V. Acknowledgements

The authors would like to thank post baccalaureate students, Mark Kellish and Jenna Peters for their invaluable efforts on data collection for this project. The authors would also like to thank Yenny Bellido Fuentes and Catya Lopez, the mid wives at La Liga Contra el Cancer, Lima Peru for their clinical assistance in data collection. This work is partially supported by NIH, NSF, and the DoD.

